# Targeting microglial P2Y12-receptor signalling attenuates neurodegeneration in experimental Parkinson’s disease

**DOI:** 10.64898/2026.10.02.756181

**Authors:** András Iring, Dóra J. Kiss, Mária Baranyi, Zsuzsanna T. Gál, Emese Herczku, Pál Tod, Attila Egyed, Kornél Demeter, Ivar von Kügelgen, Long-Jun Wu, György M. Keserű, Beáta Sperlágh

## Abstract

Neuroinflammation is increasingly recognized as a determinant of dopaminergic neuron vulnerability in Parkinson’s disease, yet the receptor-level mechanisms linking microglia to disease progression remain incompletely defined. We investigated whether microglial P2Y12 receptor (P2Y12R) signalling drives neurodegeneration in experimental Parkinsonism and assessed its pharmacological tractability. Tamoxifen-inducible microglia-specific P2ry12 deletion was examined in acute and subchronic MPTP models, with receptor deletion induced before or during disease progression. Neurodegeneration, neuroinflammation, neurochemistry, and motor function were evaluated, and novel P2Y12R antagonists identified by structure-based virtual screening were characterized in vitro and following central administration in vivo. Inducible microglia-specific P2Y12R deletion reduced nigrostriatal dopaminergic neuron loss, preserved dopamine synthesis, attenuated acute and sustained neuroinflammatory responses, and improved parkinsonian motor deficits without affecting survival. Importantly, delayed receptor deletion during ongoing neurodegeneration remained protective, indicating that P2Y12R contributes to disease progression. Two novel antagonists exhibited functional P2Y12R inhibition in vitro, and central administration of the most promising candidate recapitulated the neuroprotective phenotype observed following genetic receptor deletion. These findings identify microglial P2Y12R as a therapeutically relevant regulator of neuroinflammation and dopaminergic vulnerability and provide complementary genetic and pharmacological proof-of-concept for targeting central P2Y12R in Parkinsonism.

## 1. Introduction

Parkinson’s disease is a progressively disabling neurodegenerative disorder whose incidence and prevalence continue to rise with population aging, imposing a substantial public health burden [1]. The pathological hallmark features of Parkinson’s disease include the relatively selective loss of dopaminergic neurons in the substantia nigra pars compacta and the accumulation of misfolded α-synuclein within Lewy bodies [2]. The distinctive motor symptoms of the disease, *i.e.* asymmetric tremor, cogwheel rigidity, bradykinesia, and postural instability [3] are accompanied by a broad spectrum of nonmotor manifestations, including autonomic dysfunction and cognitive impairment as the disease progresses [4, 5]. While the precise initiating events of Parkinson’s disease remain uncertain, converging data implicate genetic susceptibility, environmental exposures, and immune dysregulation in a multifactorial model of disease pathogenesis [6]. Emerging evidence further indicates that neuroinflammatory mechanisms intersect with these pathways, positioning immune signalling as a central contributor to neuronal vulnerability rather than secondary epiphenomena [7]. Consistent with this view, human genetic and transcriptomic studies implicate microglial gene programs, including *P2RY12*, in Parkinson’s disease susceptibility. While not causal, these analyses reinforce the relevance of microglial receptor signalling pathways to disease pathogenesis [8].

The ADP-sensitive purinergic P2Y12-receptor (P2Y12R) is a Gi-coupled G protein–coupled receptor, and is the molecular target of widely prescribed antiplatelet medications used to prevent stroke and myocardial infarction [9]. Despite their clinical success, clinically approved P2Y12R inhibitors are largely restricted to the periphery and do not achieve meaningful blood-brain barrier (BBB) penetration [10]. Although brain-penetrant P2Y12R ligands have recently been developed for molecular imaging applications, therapeutic modulation of central P2Y12R signalling remains limited [11]. In the central nervous system (CNS), P2Y12R expression is highly enriched in microglia, distinguishing these cells from other myeloid populations and positioning the receptor as a potentially selective modulator of neuroimmune function [12]. Microglial P2Y12R signalling regulates essential aspects of microglial physiology, including polarization, directed migration, and process extension toward sites of neuronal injury [13]. Under physiological conditions, P2Y12R supports continuous microglial surveillance and formation of specialized microglia–neuron somatic purinergic junctions that monitor neuronal function, while also mediating directed responses to injury [14]. In addition, P2Y12R activity modulates inflammatory mediator production that influences neuronal excitability and vulnerability [15, 16]. Disruption or maladaptive engagement of these P2Y12R-dependent microglial programs may shift microglia from homeostatic surveillance toward sustained inflammatory activation, thereby contributing to neuronal vulnerability in neurodegenerative disorders such as Parkinson’s disease. In Parkinson’s disease, microglia are strongly activated by aggregated α-synuclein, neurotoxic injury, and damage-associated signals, and although acute activation may support tissue repair and debris clearance, sustained or dysregulated responses promote chronic inflammation, synaptic dysfunction, and progressive dopaminergic neurodegeneration [17–19]. Despite substantial evidence implicating microglia in Parkinson’s disease pathogenesis, the receptor-level mechanisms that shape microglial responses remain incompletely defined, constraining the development of targeted therapeutic strategies [20]. Current treatments for Parkinson’s disease largely provide symptomatic benefit through restoration of dopaminergic neurotransmission and do not engage upstream pathogenic pathways; consequently, their efficacy diminishes as neurodegeneration progresses [21]. These limitations underscore the need to identify molecular regulators of neuroinflammatory signalling and protein homeostasis.

Although molecular imaging studies have demonstrated that P2Y12R is accessible within the CNS, the functional consequences of receptor modulation during neurodegeneration have not been directly established [11]. In our previous study [18], global genetic deletion and central pharmacological inhibition of P2Y12R were both protective in MPTP-induced models of Parkinsonism; however, the specific cellular compartment mediating this effect was not defined. The present study was therefore designed to define the contribution of microglial P2Y12R by combining inducible, cell type–specific genetic approaches with the development of novel antagonists intended to act within the CNS. We hypothesized that microglial P2Y12R promotes neuroinflammatory responses that contribute to nigrostriatal degeneration in experimental Parkinsonism, and that selective genetic or pharmacological inhibition of this pathway would confer neuroprotection. Although the chemical properties of the identified compounds preclude definitive conclusions regarding passive BBB penetration, our findings identify microglial P2Y12R as a fundamental contributor of the protective action in this context, establish a platform for CNS-directed targeting, and provide a rational starting point for further pharmacological optimization.

## 2. Materials and methods

### 2.1. Animals

Animals were housed under a 12-hour light–dark cycle in a temperature- (23 ± 2°C) and humidity-controlled environment (50 ± 10%) with *ad libitum* access to food (ssniff® Souris- Elevage E, 10 mm pellet, Cat# S8189-S096; ssniff Spezialdiäten GmbH, Soest, Germany) and water. In total, five adult littermates per cage were housed in standard mouse cages with corncob bedding. Cardboard bedding materials and tubes were placed in all cages for environmental enrichment. All experiments were performed during the light phase. Studies were conducted using inducible, microglia-specific *P2ry12*-deficient mice (CX_3_CR1^CreERT2/+^- *P2ry12^fl/fl^*) and Cre-negative littermate controls (*P2ry12^fl/fl^*). Mice of both sexes, aged 8–14 weeks (27 ± 1.4 g), were used. *P2ry12^fl/fl^*mice were generated by CRISPR/Cas9-mediated insertion of loxP sites flanking exon four and were provided by Prof. Long-Jun Wu (UTHealth Houston, TX, USA). These mice were crossed with CX_3_CR1^CreERT2/+^ mice (The Jackson Laboratory, Cat. No. 025524) to obtain inducible knockout animals. All lines were backcrossed onto a C57BL/6N background for at least 8–10 generations. Cloning strategy and genotyping have been described previously [22]. All animal procedures were conducted in accordance with institutional and national guidelines (Hungarian Act 40/2013; EU Directive 2010/63/EU) and approved by the Animal Care and Experimentation Committee of the HUN-REN Institute of Experimental Medicine and the Animal Health and Food Control Station, Budapest (PE/EA/901-7/2020). Experiments complied with ARRIVE 2.0 guidelines [23, 24], and with the recommendations made by the *British Journal of Pharmacology* [24, 25]. The numbers of mice in each experimental group are shown in the corresponding figure legends. The mice were randomly assigned to experimental groups, and the investigators were blinded to the experimental status of the mice.

### 2.2. Tamoxifen induction

Cre recombinase–mediated recombination was induced by two intraperitoneal injections of tamoxifen (Sigma-Aldrich, Cat. No. T5648; 100 mg / kg) administered 48 hours apart at 4–6 weeks of age. Experiments were conducted 3–4 weeks after the final injection to allow for recombination and tamoxifen clearance. Recombination efficiency varied between animals, consistent with inducible CreERT2 systems [26, 27]. Deletion of P2Y12R was therefore validated in each cohort by immunofluorescence or immunoblotting. Animals lacking evidence of recombination in assays where verification was feasible were excluded based on predefined criteria. Across experiments, 21.8% of tamoxifen-treated animals were excluded due to absent recombination or unrelated criteria. For molecular and neurochemical analyses requiring tissue homogenization, samples were derived from cohorts with independently confirmed recombination efficiency. Two cohorts lacking detectable recombination were excluded and replaced.

### 2.3. Intracisternal administration

Mice were anesthetized with isoflurane and placed on a stereotaxic frame with temperature control. Following exposure of the cisterna magna, a glass capillary was inserted under microscopic guidance, and artificial cerebrospinal fluid (ACSF) containing P2Y12R antagonist or vehicle was injected over 1–2 minutes. The incision was sutured, and animals recovered under monitoring with postoperative analgesia (buprenorphine, 0.05 mg / kg, subcutaneous) [28]. Injections were performed 18 hours prior to MPTP administration.

### 2.4. MPTP treatment

#### 2.4.1. Acute model

Mice were randomly assigned to experimental groups and received four intraperitoneal injections of MPTP (15 mg / kg; Selleck Chemicals) at two hour intervals, as described previously [18]. Behavioural testing was performed 72 hours after the first injection. Animals were euthanized at 72 hours for assessment of microglial activation, cytokine levels, biogenic amine content (HPLC-EC), and histochemistry.

#### 2.4.2. Subchronic model

Mice received daily intraperitoneal injections of MPTP (15 mg / kg) or saline for five consecutive days (Days 1–5) [29, 30]. Tamoxifen or vehicle was administered on Day 3 and Day 5. Behavioural testing was conducted 28 days after the initial MPTP injection, and animals were euthanized 28 days after completion of treatment for histological and neurochemical analyses.

### 2.5. Behavioural analyses

#### 2.5.1. Open-field test

Locomotor activity was assessed in a 40 × 40 cm arena under dim lighting for ten minutes using EthoVision XT15 (Noldus). Total distance travelled, average velocity, and time spent in the centre zone were quantified.

#### 2.5.2. Rotarod

Motor coordination was assessed using an accelerating rotarod (IITC). Following habituation, mice were tested using fast (5–45 rpm over 30 s) and slow (1–45 rpm over 120 s) acceleration paradigms. Performance was recorded as latency to fall (maximum 180 s).

#### 2.5.3. Forelimb grip strength

Grip strength was measured using a grip strength meter (Harvard Apparatus). Mice performed five trials, and the mean of the three highest values was used for analysis.

All behavioural testing was conducted by an experimenter blinded to genotype and treatment.

### 2.6. Tyrosine hydroxylase (TH) immunohistochemistry

Mice were euthanized and transcardially perfused with PBS followed by 4 % PFA. Brains were post-fixed overnight (4 °C), sectioned coronally (40 μm), and processed as free-floating sections [31]. After quenching endogenous peroxidase activity and blocking, sections were incubated overnight with anti-TH antibody (Sigma-Aldrich, AB152, Lot. No. 3256647, RRID:AB_390204). Detection was performed using HRP-based immunohistochemistry with DAB (Vector Laboratories). Sections were mounted and cover slipped for analysis.

### 2.7. Stereological estimation

TH-positive neurons in the substantia nigra were quantified using unbiased stereology on every fourth section (ten sections total; 160 μm spacing) as described previously [31]. Neurons were identified based on morphology and TH immunoreactivity. Counting parameters included a 300 × 300 μm grid, 50 × 50 μm counting frame, and 3 μm guard zones. Total neuron number was calculated using standard stereological formulas [32], and only estimates with a coefficient of error <0.10 were included [33]. Analyses were performed blinded to group assignment. The *n* number displayed on the figures and marked in the figure legends signify the number of animals used for the indicated experimental condition.

### 2.8. Immunofluorescence and confocal imaging

Sections were processed using standard protocols and incubated with antibodies against CD68 (1:250; Bio Rad Laboratories, Cat. No. MCA1957, Lot. No. 165974, RRID:AB_322219), Iba1 (1:500; Synaptic System GmbH, Cat. No. 234004, Lot. No. 2-20, RRID:AB_2884925), and P2Y12R (1:1000; AnaSpec, Cat. No. AS 55043A, Lot. No. UB1701, RRID:AB_2298886), followed by Alexa Fluor–conjugated secondary antibodies and nuclear staining. Images were acquired using a Nikon confocal microscope under identical settings. Signal intensity was quantified in ImageJ after background subtraction, with analysis performed on multiple fields and sections per animal as described previously [34].

### 2.9. HPLC analysis of monoamines

Biogenic amines were quantified in substantia nigra and striatum by HPLC with electrochemical detection. Tissue samples were homogenized in perchloric acid containing theophylline (as an internal standard) at a concentration of 10^-5^ M and 5 × 10^-5^ M sodium metabisulfite (antioxidant of biogenic amines), centrifuged, and processed for analysis. Separation was performed using column-switching chromatography with phenyl-hexyl analytical columns, and detection was conducted at 0.73 V. Protein concentration was measured according to Lowry et al. [35]. Concentrations were calculated using an internal standard method and expressed as pmol / mg protein.

### 2.10. Cytokine measurement

Brain tissues were collected 72 hours (acute) or 28 days (subchronic) after MPTP treatment. Following homogenization in RIPA buffer with protease inhibitors and protein quantification, cytokine levels (IL-1β, IL-6, IFN-γ, TNF-α, KC, MCP-1) were measured using cytometric bead arrays (BD Biosciences) and analysed by flow cytometry [36, 37]. Values were normalized to total protein and expressed as pg / mg protein.

### 2.11. Western blotting

Brain lysates were prepared in RIPA buffer with protease inhibitors, and protein concentrations were determined by BCA assay. Equal amounts of protein were separated by SDS–PAGE, transferred to nitrocellulose membranes, and probed with antibodies against P2Y12R (1:1000; AnaSpec; Cat. No. AS 55043A, Lot. No. UB1701, RRID:AB_2298886) and GAPDH (1:3000; Cell Signaling Technology; Cat. No. 2118, Lot. No. 14, RRID:AB_561053). Bands were detected by chemiluminescence and quantified using ImageJ, with normalization to GAPDH.

### 2.12. Virtual inhibitory compound screening

Potential inhibitory compounds were identified by virtual screening of the HUN-REN RCNS compound collection against human P2Y12R using the antagonist-bound crystal structure (PDB ID: 4NTJ) [38], obtained from GPCRdb [39]. All modelling studies were carried out by the Schrödinger Small Molecule Drug Discovery Program Package [40]. The structure was prepared using Schrödinger’s Protein Preparation Wizard, missing hydrogen atoms were added, and the protonation states were assigned. An inner grid box of 12 Å x 12 Å x 12 Å was centred on the co-crystallized ligand while the outer box size was set to 48 Å x 48 Å x 48 Å. Ligand protonation states at pH 7.4 and stereoisomers were generated using LigPrep. Docking was carried out with Schrödinger’s Glide using standard precision (SP) mode. The OPLS4 force field [41] was used throughout the modelling steps. For each ligand, five poses were evaluated in post-docking minimization, and one pose was saved for visualization. No positional or interaction constraints were applied during primary screening. Hit selection was based on docking score, ligand efficiency, and visual inspection of binding poses, guided by key receptor–ligand interactions identified in the crystal structure [42].

### 2.13. Calculated CNS MPO and BBB Scores

During candidate prioritization, BBB penetration potential was estimated by calculating CNS multiparameter optimization (CNS MPO) scores according to Wager et al. [43] and BBB scores as described by Gupta et al. [44]. In addition, predicted blood–brain barrier permeability was evaluated using the LightBBB [45] and DeePred-BBB [46] computational models. Compound properties were interpreted within the scoring frameworks defined in the respective publications to estimate relative CNS penetration potential (Supplementary Table 1.).

### 2.14. Cultured cells and pharmacological assays

A CHO-K1 cell line stably expressing human P2Y12R (CHO-hP2Y12R) was used as an *in vitro* model, as previously described [47]. Briefly, CHO Flp-In cells (Invitrogen, RRID:CVCL_U424) were maintained in Ham’s F-12 medium supplemented with GlutaMAX I, 10 % foetal bovine serum, and Zeocin (100 mg / l; Invitrogen, Cat. No.: 4605059) at 36.5°C in 5 % CO_2_. Cells were transfected using LIPOFECTAMINE 2000™ (Invitrogen, Cat. No.: 11668-027), and stable clones were selected with hygromycin (500 μg / ml, Invitrogen, Cat. No.: 10687-010). The cell line was kindly provided by Prof. Ivar von Kügelgen (University of Bonn). Experiments were performed using cells at passages 13–17.

Intracellular cAMP levels were measured using a competitive enzyme immunoassay (EIA; Thermo Fisher Scientific, Cat. No. KGE002B) to assess P2Y12R activity. Cells were serum-deprived for six hours and pretreated with forskolin (10^-5^ M, Sigma Aldrich, Cat. No.: F6886; 20 min) to elevate basal cAMP levels. Three experimental conditions were applied: (i) agonist stimulation with 2-MeSADP (10^-11^ M – 10^-6^ M, Tocris, Cat. No. 1624, 10 min); (ii) preincubation with reference antagonists PSB0739 (10^-9^ M – 10^-7^ M; Tocris, Cat. No.: 3983) and ticagrelor (10^-8^ M – 10^-6^ M; Sigma Aldrich, Cat. No.: SML2482); and (iii) treatment with novel compounds (10^-5^ M), with selected candidates further tested at 10^-6^ M – 10^-5^ M. Cells were stimulated with 2-MeSADP for 10 minutes, and reactions were terminated by extraction with 0.1 M HCl. Calibration standards were used to generate a standard curve. Luminescence was measured using a Cytation 5 multi-mode plate reader (BioTek Instruments). Intracellular cAMP reduction was calculated as:

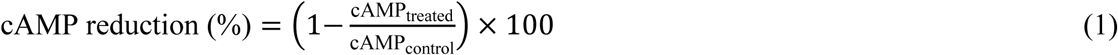

where *cAMP*_treated_ represents levels measured after agonist stimulation in the presence of antagonists, and *cAMP*_control_ corresponds to forskolin-treated cells without agonist stimulation.

### 2.15. Schild analysis

Antagonist properties were evaluated using Schild analysis based on concentration–response curves for 2-MeSADP generated in the absence and presence of fixed antagonist concentrations. Only parallel rightward shifts without reduction in maximal response were included. Dose ratios were calculated at EC_50_ and Schild plots were constructed by linear regression of log(DR − 1) versus log antagonist concentration [48, 49].

### 2.16. Cytotoxicity assay

Cytotoxicity was assessed using the CyQUANT LDH assay (Invitrogen, Cat. No.: C20300). CHO-hP2Y12R cells (6 × 10^3^ / well) were exposed to test compounds (10^-5^ M, *n* = 4) for 24 hours. LDH release was measured colorimetrically according to the manufacturer’s instructions. Cytotoxicity was calculated relative to spontaneous and maximal LDH release controls as:

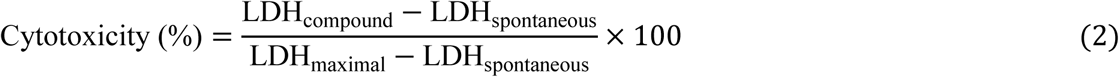

### 2.17. Experimental design and statistical analyses

Sample size was estimated using G*Power based on pilot data [50, 51]. Animals were randomly assigned to experimental groups, and all data acquisition and analyses were performed blinded. To reduce animal use, mice were reused across compatible experiments when appropriate. Exclusion criteria, including recombination failure, are detailed in Supplementary Table 2. In case of *in vivo* studies, the dose selection was based on previous studies [18, 30, 52]. Statistical analyses were performed using GraphPad Prism (v6.07). Data are presented as mean ± SD. Normality was assessed prior to analysis; parametric data were analysed using unpaired two-tailed Student’s *t* test or one-way ANOVA with Bonferroni *post hoc* correction. A *P* value <0.05 was considered significant. Experimental data and statistical analysis comply with the recommendations on experimental design and analysis in pharmacology; please see the editorial of the British Journal of Pharmacology for guidance [25].

## 3. Results

### 3.1. Microglial P2Y12-receptor deletion confers protection in experimental Parkinsonism

To investigate the role of microglial P2Y12R signalling in Parkinson’s disease–related neurodegeneration, we employed mice with inducible, microglia-specific deletion of the *P2yr12* gene (iP2Y12R-KO; iKO). Following MPTP administration, overall survival did not differ between control and iKO mice, with both groups exhibiting an approximately 35 % mortality rate (Fig. 1A), indicating that microglial P2Y12R deficiency does not influence acute lethality or systemic toxicity associated with MPTP exposure. Efficient deletion of P2Y12R was confirmed at both the protein and cellular levels. Immunoblot analysis of brain lysates revealed a marked reduction in P2Y12R protein expression in tamoxifen-treated iKO mice compared with vehicle-treated controls (Fig. 1B). Consistent with this finding, immunofluorescence analysis of the substantia nigra demonstrated a pronounced loss of P2Y12R signal in iKO mice, whereas expression of the ionized calcium-binding adapter molecule 1 (Iba1), which is known to be strongly and specifically expressed in both senescent and activated microglia [53], remained unchanged (Fig. 1C), indicating preserved microglial density despite receptor deletion. Consistent with variability inherent to inducible recombination systems, a subset of tamoxifen-treated animals exhibited incomplete receptor deletion by immunofluorescence. Animals lacking evidence of recombination were excluded from analyses where verification was feasible, and recombination efficiency was confirmed at the cohort level prior to inclusion in downstream experiments (Supplementary Table 2.).

**Figure 1.**
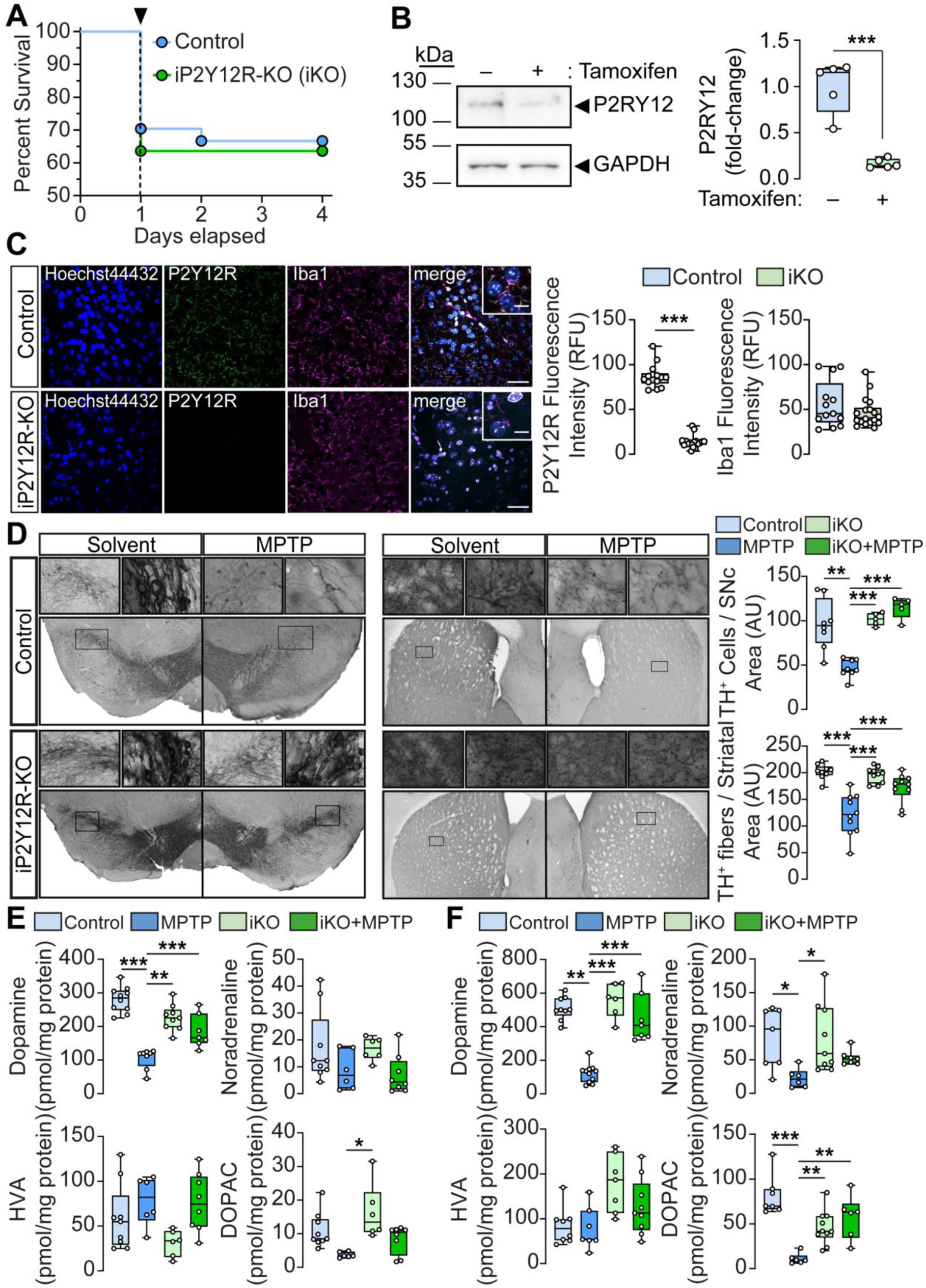
Conditional deletion of microglial P2Y12-receptors protects against MPTP- induced neurodegeneration and monoamine depletion. (A) Littermate control (Control) or inducible, microglia-specific P2Y12-receptor knockout (iKO) mice were treated with MPTP (4 × 15 mg / kg, administered at 2 hour intervals), and survival was analysed using Kaplan– Meier curves (*n* = 23 – 27). Arrow (▾) indicates the time of MPTP administration. (B) P2Y12- receptor (P2RY12) protein levels in brain lysates from Control (tamoxifen −) and iKO (tamoxifen +) mice were determined by immunoblotting. Graphs show densitometric quantification (*n* = 5). (C) Representative immunoconfocal microscopy images of brain sections from Control or iP2Y12R-KO mice, stained with antibodies directed against P2Y12R (green), Iba1 (magenta), Hoechst33342 (blue) and overlay image (merge). Scale bar: 100 µm, corresponds to 20 µm inset. Quantification of P2Y12R and Iba1 fluorescence intensity is shown in the right panel (*n* = 13 – 18). (D) Representative images of immuno-DAB staining for tyrosine hydroxylase (TH) in the substantia nigra (left panel) and striatum (right panel) following MPTP treatment in Control and iP2Y12R-KO mice. TH immunoreactivity is detected in the cell bodies and processes of dopaminergic and noradrenergic neurons. Bar graphs show quantification of TH^+^ cells in the substantia nigra pars compacta (*n* = 5 – 8) and striatum (*n* = 9 – 10). (E–F) Concentrations of dopamine, noradrenaline, homovanillic acid (HVA), and DOPAC in the substantia nigra (E) and striatum (F) of Control and iP2Y12R-KO mice 72 hours after the initial MPTP administration (*n* = 6 – 11). Box-and-whisker plots show the median, minimum and maximum values, and interquartile range. Statistical significance is indicated as \**P* ≤ 0.05, \*\**P* ≤ 0.01, \*\*\**P* ≤ 0.001 (unpaired two-tailed Student’s *t* test for B– C; one-way ANOVA with Bonferroni *post hoc* test for E–F).

We next assessed whether microglial P2Y12R deficiency modulates MPTP-induced nigrostriatal degeneration. In control mice, MPTP administration resulted in significant loss of tyrosine hydroxylase–positive (TH⁺) dopaminergic neurons in the substantia nigra pars compacta, accompanied by degenerative neuronal morphology, as well as a pronounced reduction in TH⁺ fibre density in the striatum (Fig. 1D). In contrast, iKO mice were significantly protected from MPTP-induced dopaminergic neuronal loss in the substantia nigra and from degeneration of striatal dopaminergic fibres (Fig. 1D). Neurochemical analyses corroborated these histological findings. In control mice, MPTP treatment led to a significant reduction in dopamine levels in the substantia nigra, whereas levels of noradrenaline, homovanillic acid (HVA), and DOPAC were largely preserved (Fig. 1E). In the striatum, MPTP caused pronounced depletion of dopamine, noradrenaline, and DOPAC (Fig. 1F). Notably, microglial P2Y12R deletion significantly attenuated MPTP-induced dopamine loss in the substantia nigra (Fig. 1E) and prevented striatal dopamine and DOPAC depletion (Fig. 1F). In contrast, P2Y12R deficiency did not significantly alter nigral noradrenaline, HVA, or DOPAC concentrations, nor striatal noradrenaline or HVA levels, under either basal or MPTP-treated conditions (Fig. 1E–F). Collectively, these data demonstrate that microglial P2Y12R signalling contributes to MPTP-induced nigrostriatal dopaminergic neurodegeneration and associated monoamine depletion, while not affecting acute mortality or microglial abundance.

To define the contribution of microglial P2Y12R to the acute neuroinflammatory response elicited by MPTP we examined markers of microglial activation and cytokine expression 72 hours after MPTP administration. In substantia nigra of control mice, MPTP triggered a significant increase in the fluorescence intensity of the myeloid specific lysosomal-associated membrane protein, CD68 [54], compared with vehicle-treated animals, indicating enhanced microglial activation (Fig. 2A). In contrast, immunoreactivity for P2Y12R and Iba1 was unaltered at this time point (Fig. 2A). Consistent with microglial activation, MPTP- administration induced a pronounced inflammatory response, as evidenced by significantly elevated levels of IL-1β, IL-6, IFN- γ, TNFα, Keratinocyte-Derived Chemokine (KC) and Monocyte Chemoattractant Protein-1 (MCP-1) in both the substantia nigra and the striatum (Fig. 2B and C). Conditional deletion of P2Y12R in microglia effectively abolished the MPTP-induced increase in CD68 fluorescence intensity in the substantia nigra (Fig. 2A) and attenuated the elevation of inflammatory cytokines in both brain regions examined (Fig. 2B and C). These data indicate that loss of microglial P2Y12R markedly diminishes acute MPTP-induced microglial activation and the associated proinflammatory response.

**Figure 2.**
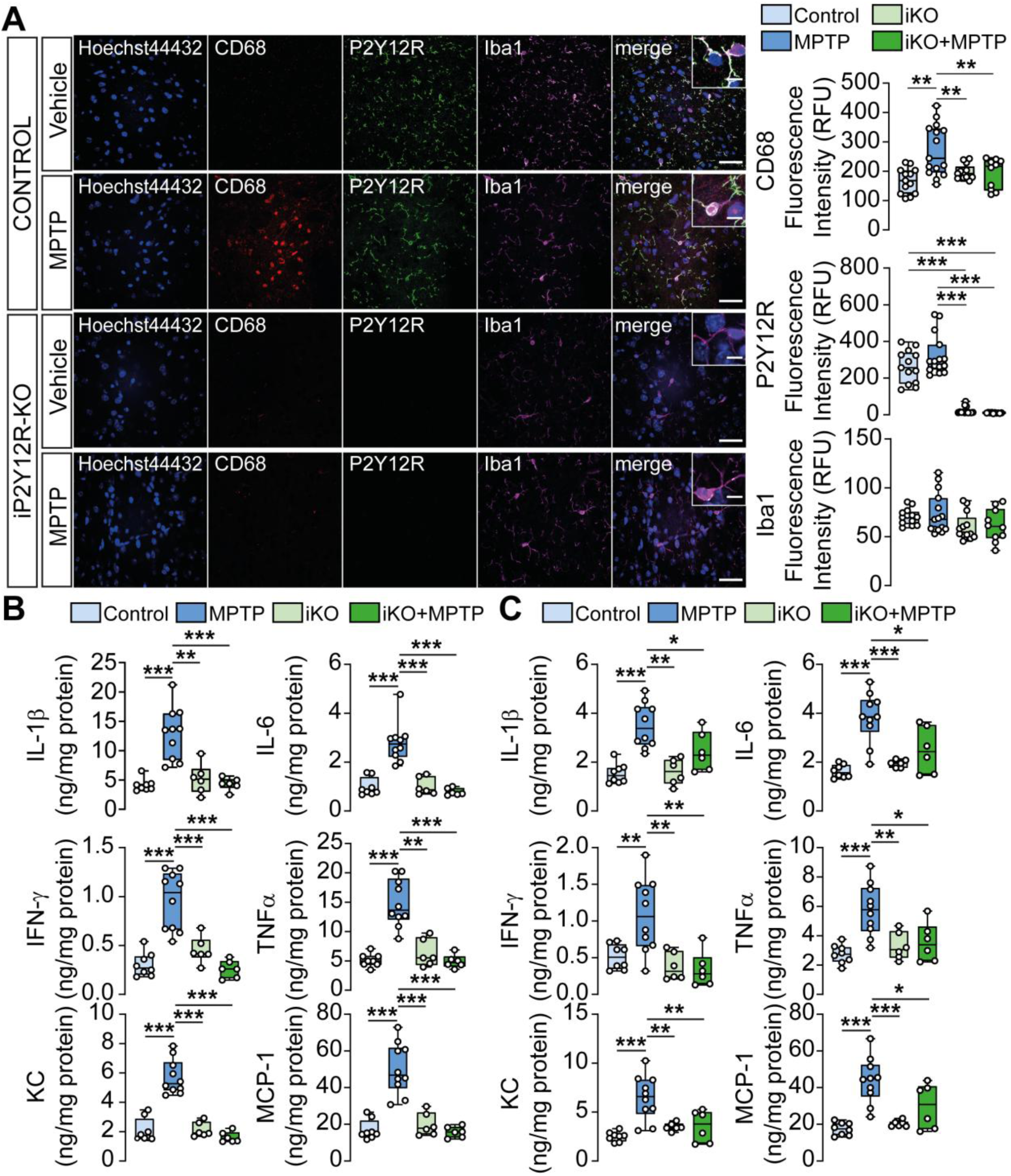
Genetic deletion of microglial P2Y12R attenuates MPTP-induced neuroinflammation and suppresses pro-inflammatory cytokine production. (A–C) Control and inducible P2Y12R knockout (iP2Y12R-KO; iKO) mice were treated with MPTP (4 × 15 mg / kg, 2 hour intervals) or vehicle. (A) Representative confocal images of brain sections from control and iKO mice stained for CD68 (red), P2Y12R (green), Iba1 (magenta), Hoechst33342 (blue) and overlay image (merge). Scale bar: 100 µm, corresponds to 20 µm inset (left panel). Quantification of CD68 fluorescence intensity (upper-right panel), P2Y12R fluorescence intensity (middle-left panel) and Iba1 fluorescence intensity (lower-right panel) (n = 10 – 15); **(B-C)** IL-1β, IL-6, IFN- γ, TNFα, Keratinocyte-Derived Chemokine (KC) and Monocyte Chemoattractant Protein-1 (MCP-1) concentration measured from substantia nigra (*n* = 6 - 8) (B) or striatum samples (*n* = 6 - 8) (C). Box-and-whisker plots show the median, minimum and maximum values, and interquartile range. Statistical significance is indicated as \**P* ≤ 0.05, \*\**P* ≤ 0.01, \*\*\**P* ≤ 0.001 (one-way ANOVA with Bonferroni *post hoc* test).

We next assessed the impact of microglial P2Y12R deletion on motor performance in the acute MPTP model. In the open field test, MPTP treatment had minimal effects on general locomotor activity, as reflected by comparable movement velocity and total distance travelled across experimental groups (Fig. 3A). Likewise, no significant differences were observed in the time spent in the centre of the arena, indicating preserved exploratory behaviour and absence of overt anxiety-like or cognitive deficits (Fig. 3A). Despite preserved general locomotion, MPTP-treated mice exhibited clear motor coordination impairments. Performance on the accelerating rotarod revealed a significant reduction in latency to fall under both fast and slow ramp-up protocols, consistent with akinetic and bradykinetic movement patterns as well as gait abnormalities (Fig. 3B). In addition, MPTP administration increased forelimb grip force, reflecting muscle rigidity and impaired muscle relaxation rather than enhanced muscle strength (Fig. 3C). Microglial P2Y12R deletion did not alter baseline locomotor activity or exploratory behaviour but significantly ameliorated MPTP-induced motor deficits. P2Y12R-deficient mice demonstrated improved performance on both rotarod protocols and a distinct reduction in muscle rigidity, as evidenced by normalized grip strength following MPTP treatment (Fig. 3A– C). These results demonstrate that loss of microglial P2Y12R confers protection against MPTP-induced parkinsonian motor deficits.

**Figure 3.**
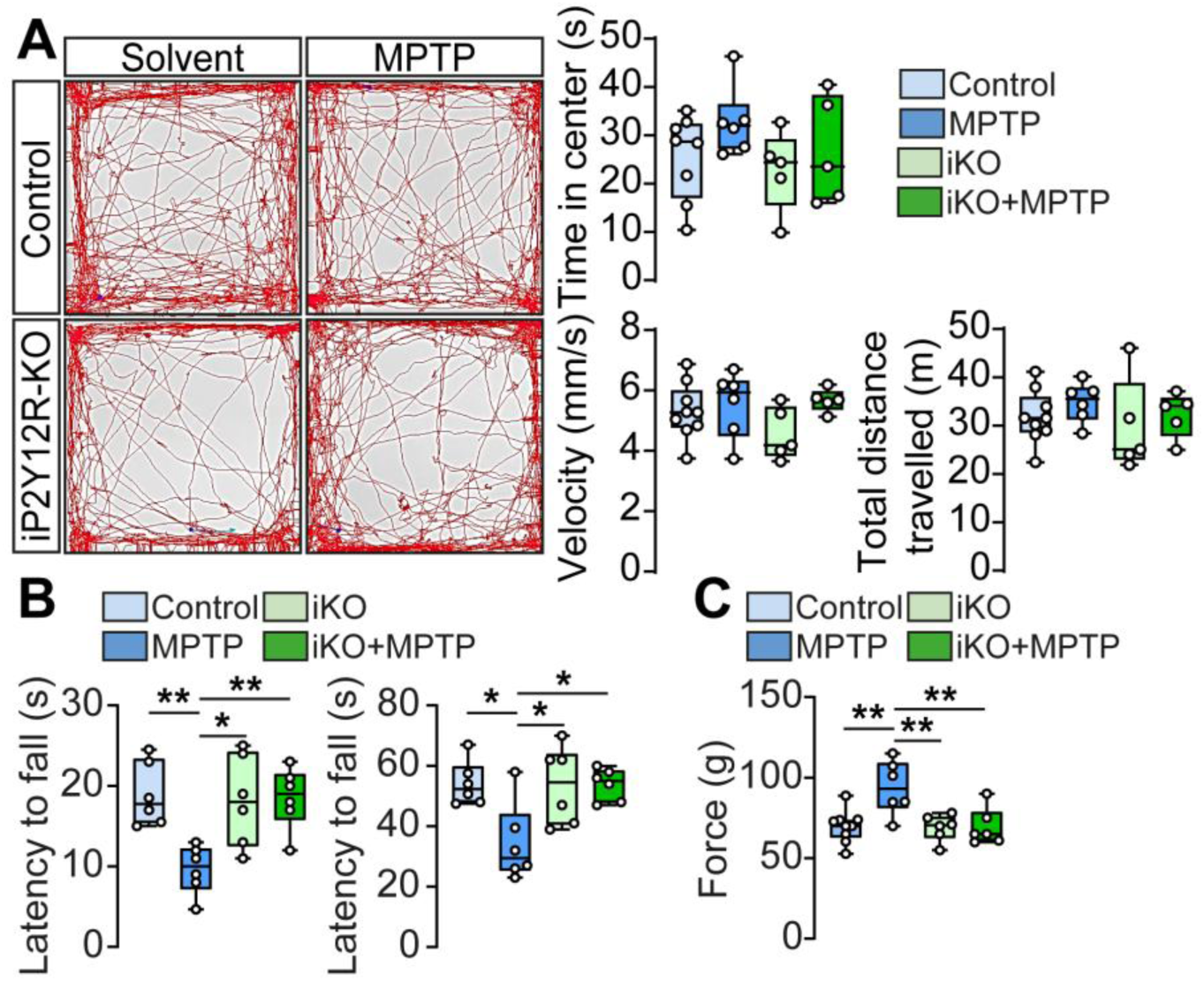
Microglial P2Y12R deletion ameliorates MPTP-induced Parkinsonian motor deficits without affecting spontaneous locomotion. **(A)** Representative open-field trajectories of Control and iP2Y12R-KO (iKO) mice treated with vehicle or MPTP (4 × 15 mg / kg, 2 hour intervals). Quantification of time spent in the centre, mean velocity, and total distance travelled is shown on the right (*n* = 5 - 8). **(B)** Motor coordination and balance assessed using the rotarod test. The left panel shows latency to fall during a fast accelerating protocol, sensitive to akinesia and bradykinesia, while the right panel shows performance during a slow accelerating protocol, reflecting general motor coordination and balance (*n* = 6). **(C)** Forelimb grip strength measured as maximal force (*n* = 6 - 9). Data are presented as box-and-whisker plots, displaying the median, minimum and maximum values, and interquartile range. Statistical significance is indicated as \**P* ≤ 0.05, \*\**P* ≤ 0.01 (one-way ANOVA with Bonferroni *post hoc* test).

### 3.2. Targeting microglial P2Y12R during ongoing neurodegeneration confers neuroprotection

To determine whether targeting microglial P2Y12R signalling during ongoing neurodegeneration alters disease progression, we used a subchronic MPTP model in which conditional recombination was induced after initiation of nigrostriatal injury. Mice received daily MPTP injections for five consecutive days, with tamoxifen administered on treatment Days 3 and 5 to trigger microglia-specific deletion of P2Y12R. Nigrostriatal integrity was evaluated 28 days after the initial MPTP administration. In control mice, subchronic MPTP administration resulted in a pronounced loss of TH⁺ dopaminergic neurons in the substantia nigra pars compacta (SNpc), accompanied by a pronounced depletion of striatal TH⁺ dopaminergic innervation (Fig. 4A). This structural degeneration was paralleled by significant reduction in dopamine and noradrenaline levels in the substantia nigra (Fig. 4B), as well as decreased concentrations of dopamine, noradrenaline, and the dopamine metabolite DOPAC in the striatum (Fig. 4C). In contrast, mice with inducible microglial P2Y12R deletion were significantly protected from MPTP-induced nigrostriatal degeneration, exhibiting preserved TH⁺ neuronal numbers in the SNpc, reduced loss of striatal dopaminergic fibres (Fig. 4A), and retention of monoamine levels in both the substantia nigra and striatum (Fig. 4B and C).

**Figure 4.**
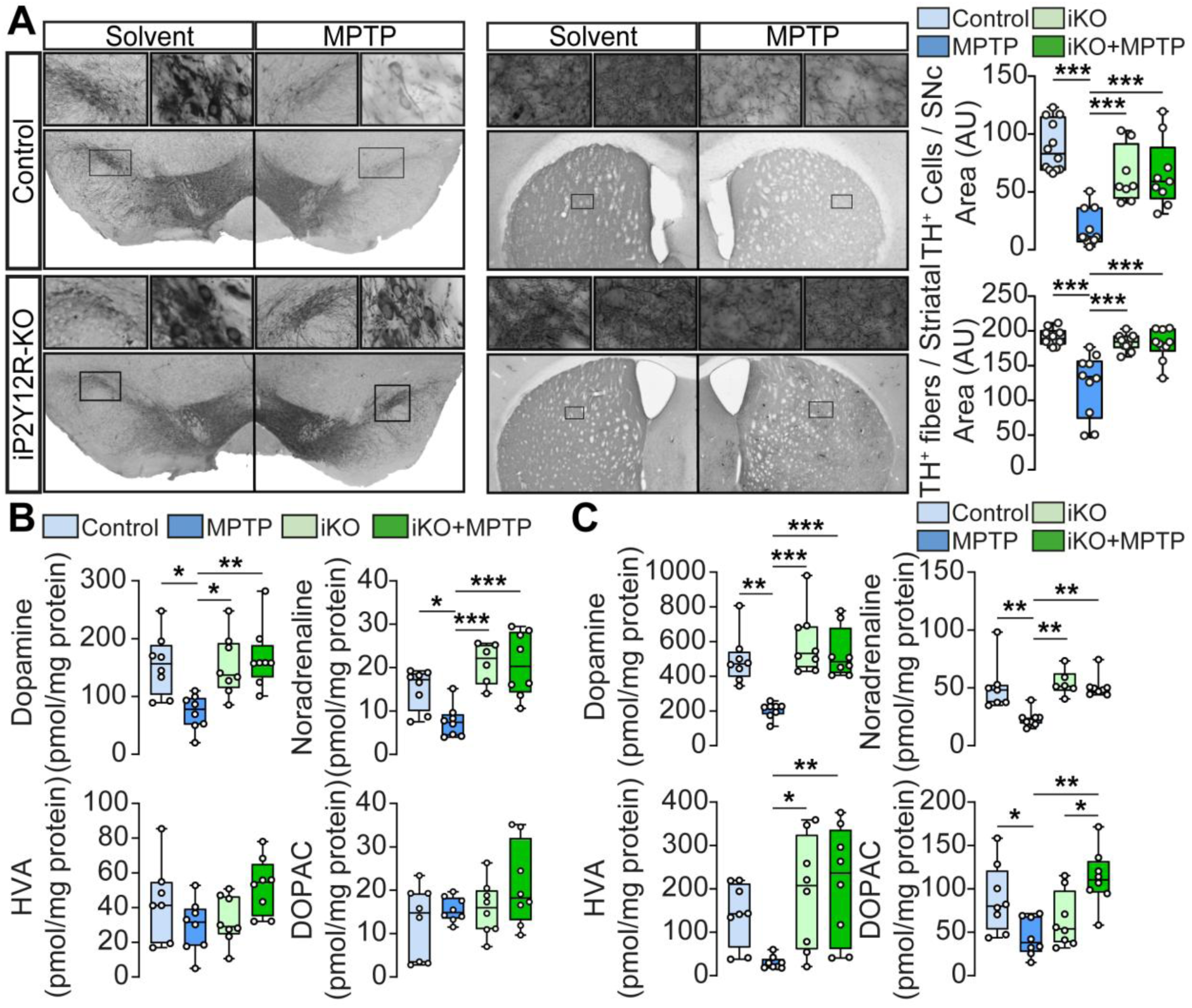
Temporal blockade of microglial P2Y12R halts progression of subchronic Parkinsonism. **(A-C)** Control and inducible, microglia-specific P2Y12-receptor knockout (iP2Y12R-KO; iKO) mice were treated with MPTP (15 mg / kg, once daily for five consecutive days). To trigger microglia-specific recombination during evolving pathology, iKO mice received tamoxifen on Days 3 and 5 of MPTP treatment. All histological and neurochemical analyses were performed four weeks after initiation of the MPTP regimen. **(A)** Representative images of immuno-DAB staining for TH in the substantia nigra (left panel) and striatum (right panel). TH immunoreactivity is detected in the cell bodies and processes of dopaminergic and noradrenergic neurons. Bar graphs show quantification of TH^+^ cells in the substantia nigra pars compacta (*n* = 8 – 12) and striatum (*n* = 9 – 10). **(B–C)** Concentrations of dopamine, noradrenaline, homovanillic acid (HVA), and DOPAC in the substantia nigra (B) and striatum (C) of Control and iP2Y12R-KO mice (*n* = 6 – 8). Box-and-whisker plots show the median, minimum and maximum values, and interquartile range. Statistical significance is indicated as \**P* ≤ 0.05, \*\**P* ≤ 0.01, \*\*\**P* ≤ 0.001 (one-way ANOVA with Bonferroni *post hoc* test).

Given the established role of microglial activation in sustaining chronic neurodegeneration in MPTP-induced parkinsonism, we next examined microglial activation and cytokine production during the subchronic phase of disease progression. In control mice, MPTP induced a significant increase in CD68 fluorescence intensity, consistent with sustained microglial activation, whereas immunoreactivity for P2Y12R and the pan-microglial marker Iba1 remained unchanged (Fig. 5A). Conditional deletion of microglial P2Y12R during disease progression attenuated the MPTP-induced increase in CD68 fluorescence intensity, without affecting Iba1 levels (Fig. 5A), indicating a selective suppression of the activated microglial phenotype. In contrast to the acute neurotoxic regimen, which elicited inflammatory responses in both the SNpc and striatum, the subchronic MPTP paradigm was associated with a more spatially restricted inflammatory profile, characterized by increased TNFα and IL-1β production in the striatum (Fig. 5C) but not in the SNpc (Fig. 5B). These findings suggest that microglial P2Y12R signalling contributes to the maintenance of a neuroinflammatory milieu during subchronic parkinsonism.

**Figure 5.**
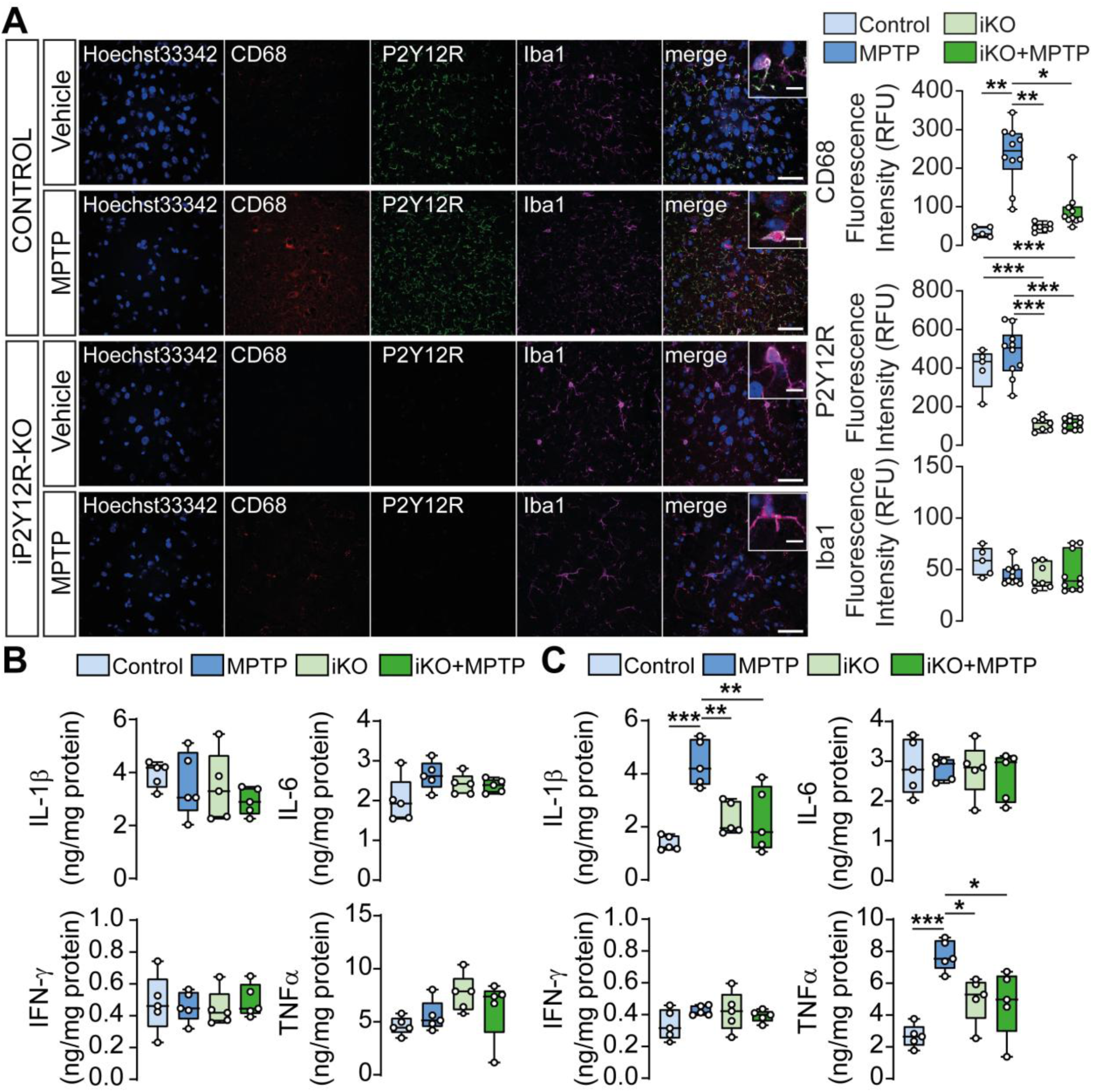
Inducible microglial P2Y12R deletion restrains inflammatory microglial responses in subchronic Parkinsonism. **(A-C)** Control and inducible microglia-specific P2Y12-receptor knockout (iP2Y12R-KO; iKO) mice were subjected to the subchronic MPTP regimen and tamoxifen-induced recombination. All analyses were performed four weeks after initiation of MPTP treatment. **(A)** Representative confocal images of brain sections stained for CD68 (red), P2Y12R (green), Iba1 (magenta), Hoechst33342 (blue) and overlay image (merge). Scale bar: 100 µm, corresponds to 20 µm inset (left panel). Quantification of CD68 fluorescence intensity (upper-right panel), P2Y12R fluorescence intensity (middle-left panel) and Iba1 fluorescence intensity (lower-right panel) (*n* = 5 – 10). **(B-C)** IL-1β, IL-6, IFN- γ, TNFα concentration measured from substantia nigra (*n* = 5) (B) or striatum samples (*n* = 5) (C). Box-and-whisker plots show the median, minimum and maximum values, and interquartile range. Statistical significance is indicated as \**P* ≤ 0.05, \*\**P* ≤ 0.01, \*\*\**P* ≤ 0.001 (one-way ANOVA with Bonferroni *post hoc* test).

We next examined whether preservation of nigrostriatal integrity following microglial P2Y12R deletion translated into functional improvement. In the open-field test, subchronic MPTP treatment had minimal effects on time spent in the centre, locomotor velocity, or total distance travelled across experimental groups (Fig. 6A), indicating preserved baseline locomotion and exploratory behaviour. In the accelerating rotarod assay, MPTP-treated control mice exhibited a significant reduction in latency to fall under the fast ramp-up protocol, but not under the slow ramp-up protocol, indicating a selective impairment in motor coordination and bradykinetic movement patterns, whereas generalized gait abnormalities were not a prominent feature of the subchronic model (Fig. 6B). Additionally, subchronic MPTP administration induced increased forelimb grip force, reflecting muscle rigidity (Fig. 6C). Inducible loss of microglial P2Y12R did not affect baseline motor performance but significantly ameliorated MPTP-induced motor deficits and reduced muscle rigidity (Fig. 6A–C).

**Figure 6.**
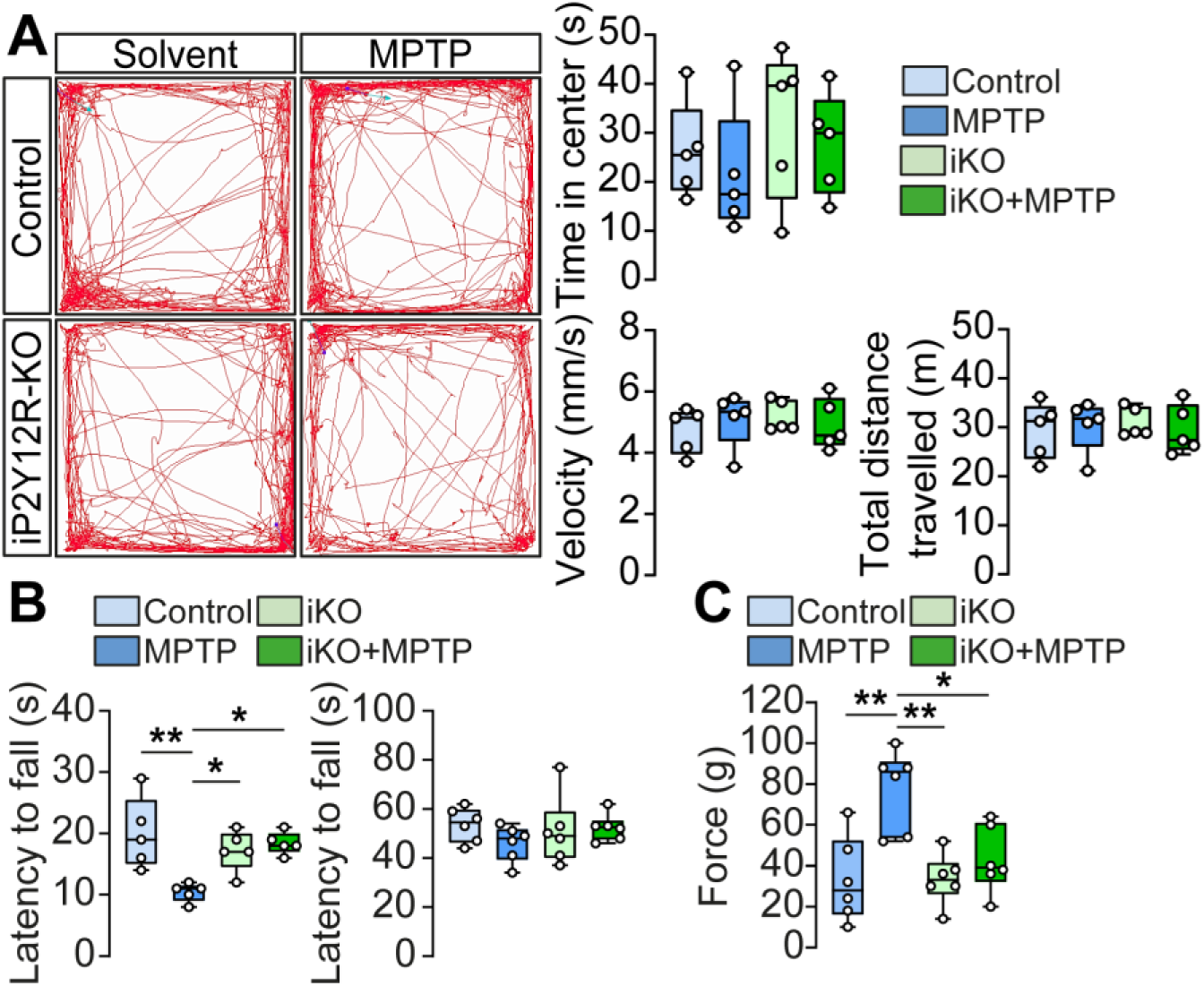
Microglial P2Y12R deletion improves motor function in subchronic Parkinsonism. **(A–C)** Control and inducible microglia-specific P2Y12-receptor knockout (iP2Y12R-KO; iKO) mice received MPTP (15 mg / kg, once daily for 5 days). To induce microglial recombination during ongoing pathology, iKO mice were treated with tamoxifen on Days 3 and 5. All analyses were performed four weeks after initiation of MPTP treatment. **(A)** Representative open-field trajectories of Control and iKO mice treated with solvent or MPTP. Quantification of time spent in the centre, mean velocity, and total distance travelled is shown (*n* = 5). **(B)** Motor coordination and balance assessed by the rotarod test. The left panel shows performance during a fast accelerating protocol, sensitive to akinesia and bradykinesia, while the right panel shows performance during a slow accelerating protocol, reflecting general motor coordination and balance (*n* = 5 – 6). **(C)** Forelimb grip strength measured as maximal force (*n* = 6). Data are presented as box-and-whisker plots, displaying the median, minimum and maximum values, and interquartile range. Statistical significance is indicated as \**P* ≤ 0.05, \*\**P* ≤ 0.01 (one-way ANOVA with Bonferroni *post hoc* test).

Collectively, these results demonstrate that acute, conditional loss of microglial P2Y12R signalling during ongoing neurodegeneration attenuates sustained neuroinflammation, preserves functional dopaminergic neurons, and mitigates parkinsonian motor deficits, thereby strengthening the rationale for the development of CNS-directed P2Y12R inhibitors.

### 3.3. Development and characterization of neuroprotective P2Y12R inhibitors

Building on this mechanistic paradigm and prompted by the limited brain penetration of clinically approved P2Y12R antagonists, we sought to develop compounds capable of engaging central P2Y12R signalling. Virtual screening of the HUN-REN RCNS chemical library against the antagonist-bound human P2Y12R identified candidate ligands, predominantly N-substituted phenothiazines and related heterocycles, with predicted binding affinity. Selected hits were subsequently evaluated using classical pharmacological approaches in CHO-K1 cells stably expressing human P2Y12R. Antagonist potency was quantified using dose-ratio analysis and Schild regression to derive pA_2_ values [49]. Because P2Y12R is Gi-coupled receptor, intracellular cAMP accumulation was stimulated with forskolin, and receptor activation was assessed as inhibition of forskolin-induced cAMP by the selective P2Y12R agonist 2-MeSADP [42]. Assay performance was validated using the well-characterized competitive antagonist PSB0739 and the clinically approved reversible allosteric modulator ticagrelor [55, 56]. Both compounds produced concentration-dependent rightward shifts of the 2-MeSADP concentration–response curve (Fig. 7A and B). Schild regression yielded slopes close to unity; for ticagrelor, this behaviour is consistent with surmountable allosteric modulation rather than classical competitive antagonism. Importantly, these assays quantify functional antagonism at the receptor level but do not distinguish between orthosteric, allosteric, or probe-dependent mechanisms of inhibition. Derived pA_2_ values were in agreement with published reports, supporting assay robustness [47, 57].

**Figure 7.**
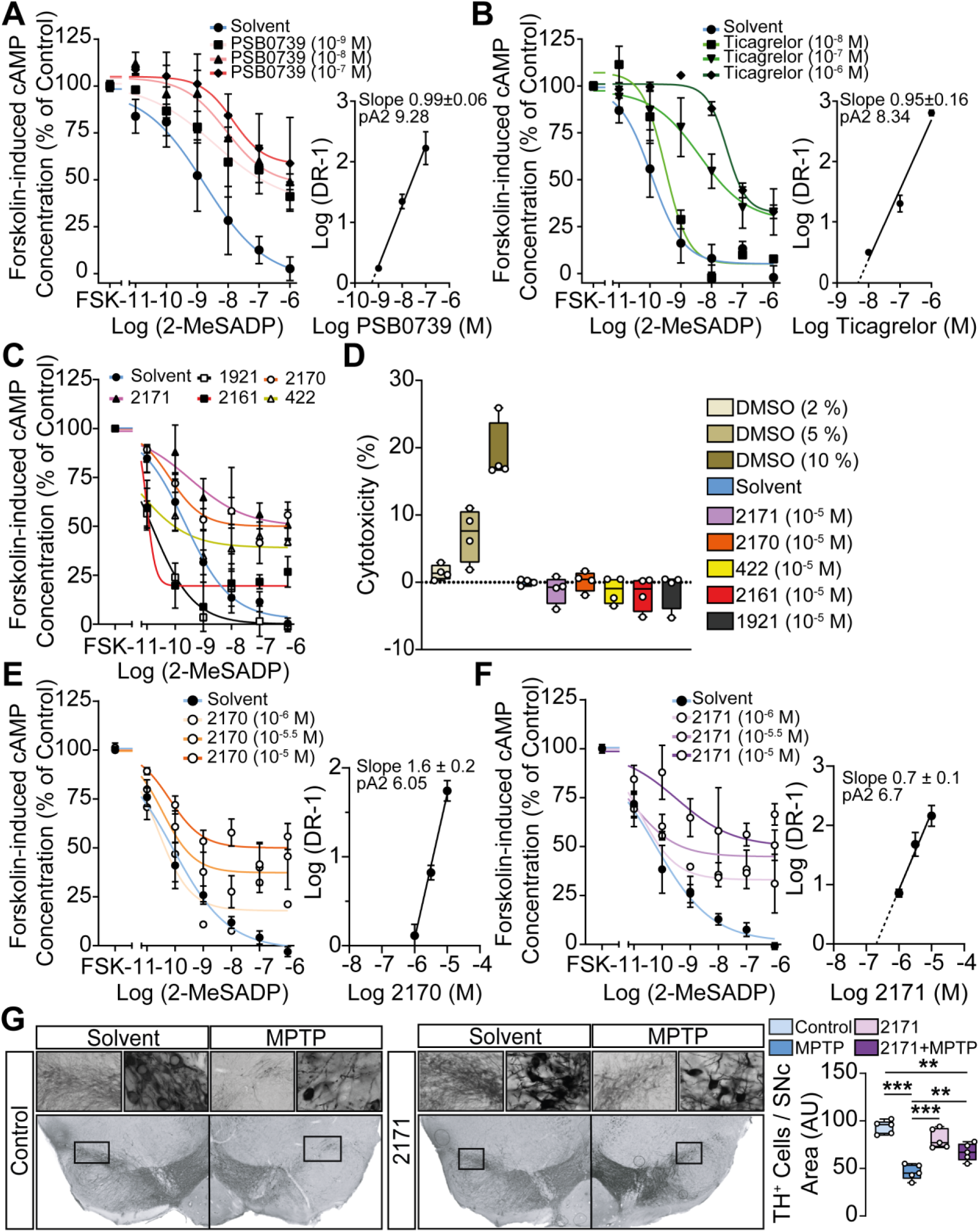
Novel P2Y12R modulators protect against Parkinsonian degeneration. **(A-B)** Antagonist activity using reference antagonists at the human P2Y12R was evaluated by measuring forskolin-induced cAMP accumulation in CHO Flp-In cells stably expressing human P2Y12R (CHO-hP2Y12R). Concentration–response curves depict forskolin-induced cAMP levels in cells stimulated with increasing concentrations of 2-MeSADP in the absence (Solvent) or presence of PSB0739 (A; 10^−9^−10^−7^ M) or ticagrelor (B; 10^−8^–10^−6^ M). cAMP levels are expressed as percentage of the forskolin response in control conditions. Corresponding Schild plots (right panels) show log (DR – 1) as a function of antagonist concentration, with calculated slopes and pA_2_ values indicated. Data are shown as mean ± SD from 5 to 20 independent experiments. **(C)** Primary testing of five novel compounds (1921, 2170, 2171, 2161, and 422; each at 10^-5^ M) for P2Y12R antagonistic activity in the CHO-hP2Y12R cells using the cAMP accumulation assay. Data are expressed as percentage of the forskolin-induced cAMP response. Data are shown as mean ± SD from 8 to 24 independent experiments. **(D)** Cytotoxicity assessment of test compounds, where cytotoxicity is presented as percentage change relative to solvent-treated cells (baseline). Cells were exposed for 24 hours to DMSO (2 %, 5 %, or 10 %) or to the indicated compounds. **(E-F)** Detailed pharmacological characterization of the two selected hit compounds, 2170 (E) and 2171 (F). Concentration-dependent antagonism of 2-MeSADP responses was assessed at multiple antagonist concentrations for 2170 (E; 10^−6^−10^−5^ M) and 2171 (F; 10^−6^−10^−5^ M). Corresponding Schild analyses (right panels) show log (DR – 1) plotted against antagonist concentration, with calculated slopes and pA_2_ values indicated. Data are shown as mean ± SD from 7 to 26 independent experiments. **(G)** Representative images of immuno-DAB staining for TH in the substantia nigra. TH immunoreactivity is detected in the cell bodies and processes of dopaminergic and noradrenergic neurons. Bar graphs show quantification of TH^+^ in the substantia nigra pars compacta (*n* = 5). Box-and-whisker plots show the median, minimum and maximum values, and interquartile range. Statistical significance is indicated as \*\**P* ≤ 0.01, \*\*\**P* ≤ 0.001 (one-way ANOVA with Bonferroni *post hoc* test).

Five compounds identified by structure-guided screening were subsequently tested at a screening concentration of 10^-5^ M (Fig. 7C). Of these, only compounds 2170 and 2171 produced a clear rightward displacement of the agonist concentration–response curve, consistent with functional antagonism at P2Y12R. To exclude nonspecific effects, cytotoxicity was assessed following 24 hour exposure to 10^-5^ M compound. None of the tested molecules significantly affected cell viability, indicating that the observed pharmacological effects were not attributable to cytotoxicity (Fig. 7D).

Further pharmacological characterization revealed that increasing concentrations of compound 2170 produced parallel rightward shifts of the 2-MeSADP concentration–response curve, consistent with inhibition of P2Y12R-mediated signalling and yielding an apparent pA₂ estimate of 6.05 (Fig. 7E). Compound 2171 displayed slightly higher apparent affinity, with an apparent pA₂ estimate of 6.7, and similarly induced concentration-dependent shifts indicative of receptor antagonism (Fig. 7F).

To evaluate neuroprotective efficacy, compound 2171 was administered into the cisterna magna (∼10^-6^ M) prior to acute MPTP challenge. MPTP induced a marked loss TH⁺ neurons in the SNpc, whereas treatment with 2171 significantly preserved TH⁺ neuronal area compared with vehicle-treated controls (Fig. 7G). Although pharmacological inhibition likely engages overlapping but non-identical receptor signalling states relative to genetic deletion, these data provide orthogonal support for P2Y12R involvement and indicate that acute central inhibition with a novel small-molecule antagonist can attenuate dopaminergic neurodegeneration. Owing to the limited solubility of compound 2171, PAMPA-based assessment of passive permeability was non-informative and did not permit conclusions regarding BBB penetration (data not shown) [58]. Therefore, CNS penetration was predicted using CNS MPO and BBB metrics. Compound 2171 achieved the highest BBB Score (4.07), consistent with predicted blood–brain barrier permeability, and a CNS MPO score of 3.95, approaching the generally accepted threshold (Supplementary Table 1.). Ongoing medicinal chemistry efforts are focused on optimizing receptor affinity and physicochemical properties to enable future evaluation of central bioavailability.

Together, these findings demonstrate that acute pharmacological inhibition of central P2Y12R mitigates nigrostriatal degeneration and identify a novel class of P2Y12R antagonists with potential for central nervous system targeting as candidate chemical starting points for CNS-directed P2Y12R modulation for Parkinson’s disease.

## 4. Discussion

The present study identifies microglial P2Y12R signalling as a functionally relevant regulator of neuroinflammation, dopaminergic vulnerability, and motor dysfunction in experimental Parkinsonism, and extends prior conceptual work by combining cell-type specific transgenic technology with first-generation efforts toward central pharmacological targeting [18]. Importantly, these findings advance P2Y12R from a mechanistically implicated microglial marker to a receptor-level pathway with demonstrable impact on disease-relevant outcomes.

A notable distinction from our earlier studies is that inducible, microglia-specific deletion of P2Y12R did not influence overall survival following acute MPTP intoxication, in contrast to the increased mortality observed after global P2Y12R deficiency or acute central pharmacological blockade [18]. This divergence indicates that survival effects attributed to P2Y12R inhibition are not solely mediated by microglial signalling but instead reflect differences in cellular scope, temporal context, and systemic vulnerability. Consistent with the preferential targeting achieved by the inducible recombination strategy, immunoblotting of whole-brain lysates from inducible knockout mice revealed a markedly reduced but detectable residual P2Y12R signal, likely reflecting P2Y12R expressed by non-microglial populations, including CNS-associated macrophages and other cells not targeted by the deletion [59, 60]. Global or centrally administered P2Y12R inhibition disrupts receptor signalling across these compartments, potentially compromising coordinated neuroimmune and neurovascular responses required for resilience to acute toxic stress [17].

Although P2Y12R expression is not exclusively restricted to microglia, the inducible Cx3cr1- CreERT2 strategy provides preferential targeting of long-lived CNS-resident microglia [61]. Following tamoxifen administration, recombination initially occurs in CX3CR1-expressing myeloid populations [61]; however, because microglia are self-renewing and persist throughout adult life, whereas circulating monocytes undergo continuous turnover, recombined peripheral myeloid cells are progressively replaced by non-recombined progenitors derived from the bone marrow [62, 63]. Consequently, after an appropriate washout period, recombination is largely retained within resident microglia, providing a widely adopted approach for microglia-selective genetic manipulation in the adult CNS. Thus, while contributions from other long-lived CX3CR1-expressing populations cannot be formally excluded, the observed phenotype is mostly consistent with a predominant role for microglial P2Y12R signalling. In contrast to global receptor deficiency, selective deletion of microglial P2Y12R dissociates lethality from neurodegeneration, preserving essential systemic and vascular functions while isolating disease-relevant neuroinflammatory mechanisms. Together, these findings suggest that P2Y12R signalling fulfils context-dependent protective functions during acute toxic stress and that broad or temporally indiscriminate inhibition may disrupt adaptive systemic or neurovascular responses.

Additionally, incomplete recombination in a subset of tamoxifen-treated animals represents an inherent limitation of inducible genetic approaches and may contribute to biological variability across cohorts [26]. However, such variability would be expected to bias results toward underestimation of the phenotypic impact of P2Y12R deletion rather than generating spurious protective effects, because non-recombined cells retain receptor function. Cohort-level validation of recombination efficiency across independent experiments supports the robustness of the observed neuroprotective phenotype despite this variability.

Within this framework, microglial P2Y12R deletion conferred substantial protection against MPTP-induced loss of nigrostriatal dopaminergic neurons and preservation of dopaminergic synthetic capacity, reflected by maintained TH⁺ neuronal counts, striatal fibre density, and monoamine levels. These findings indicate that P2Y12R signalling contributes functionally to neuronal vulnerability rather than merely reflecting a secondary marker of microglial activation. Importantly, this neuroprotection was observed without altering microglial abundance, implicating receptor-dependent functional reprogramming [64].

P2Y12R is widely regarded as a marker of homeostatic or non-inflammatory microglia, and its expression is typically reduced during the transition toward activated inflammatory states [65, 66]. In human brain, P2Y12R is enriched in parenchymal microglia and is reduced in lesion-associated myeloid cells in neuroinflammatory diseases, supporting its use as a marker of the homeostatic microglial state [65]. However, reduced expression should not be equated with complete receptor loss or functional irrelevance. Residual P2Y12R expression can nevertheless be detected in microglia during pathological states [65], and extracellular nucleotide signalling represents an important mechanism by which injured tissue communicates with microglia [66]. In particular, ATP released following cellular stress or injury can undergo extracellular conversion to ADP, providing a potential endogenous stimulus for P2Y12R [67]. Our finding that P2Y12R expression was not abolished following MPTP exposure is therefore compatible with a model in which the remaining receptor pool could continue to sense extracellular nucleotides and contribute to microglial responses during neurodegeneration. Importantly, the protective phenotype produced by inducible microglia-specific P2Y12R deletion indicates that the functional activity of this residual receptor population is sufficient to influence disease-related neuroinflammation and dopaminergic vulnerability. Thus, P2Y12R downregulation during microglial activation may represent a shift in receptor abundance rather than complete cessation of P2Y12R-dependent signalling.

Although toxin-based models do not fully recapitulate the progressive and multifactorial nature of Parkinson’s disease, they remain valuable experimental systems for dissecting mechanisms that influence dopaminergic neuronal vulnerability. Among these models, MPTP reproduces pathological features of nigrostriatal degeneration, including selective loss of dopaminergic neurons, depletion of striatal dopamine, and robust microglial activation and neuroinflammatory responses within the nigrostriatal system [68]. In comparative studies, MPTP has been shown to produce more pronounced dopaminergic neurodegeneration and neuroinflammation than chronic rotenone exposure, whereas rotenone more closely models mitochondrial complex I dysfunction and can induce α-synuclein-positive inclusions resembling Lewy body pathology [68]. Consequently, the relative strengths of these toxin-based models depend on the biological question under investigation.

The present study was designed to interrogate the contribution of microglial P2Y12R signalling to neuroinflammatory processes that influence dopaminergic vulnerability. Within this context, the robust and reproducible inflammatory phenotype of the MPTP model provides a particularly suitable platform for mechanistic investigation. Nevertheless, toxin-induced injury does not capture the full spectrum of pathological processes implicated in Parkinson’s disease, including progressive α-synuclein aggregation, proteostatic dysfunction, and the complex genetic architecture that contributes to disease susceptibility [69]. Conversely, genetic and α- synuclein-based models capture complementary dimensions of disease biology but are also associated with important limitations. Many genetic models exhibit limited nigrostriatal neurodegeneration despite clear molecular pathology, whereas α-synuclein overexpression and seeding paradigms often display variability in the extent, anatomical distribution, and temporal progression of pathology [70, 71]. Furthermore, disease progression may require prolonged observation periods and can be influenced by factors such as transgene expression levels, injection paradigms, genetic background, and experimental conditions, potentially affecting reproducibility across laboratories.

In contrast, toxin-based models provide highly reproducible and temporally controlled induction of dopaminergic injury, facilitating mechanistic interrogation of pathways that influence neuronal vulnerability and neuroinflammatory responses. Thus, although no single experimental model fully recapitulates the complexity of Parkinson’s disease, these approaches should be viewed as complementary rather than competing systems. Future studies extending the present findings to α-synuclein-based and Parkinson’s disease–associated genetic models will be important to determine whether microglial P2Y12R signalling similarly contributes to neurodegeneration in more chronic and etiologically relevant disease contexts.

Mechanistically, loss of microglial P2Y12R was associated with distinct attenuation of both acute and sustained neuroinflammatory responses to MPTP, as reflected by reduced CD68 immunoreactivity and broadly diminished induction of proinflammatory cytokines. Consistent with our earlier pharmacological observations using central P2Y12R blockade [18], these findings support a role for P2Y12R signalling in sustaining activated microglial states during dopaminergic injury, particularly across the transition from acute insult to progressive degeneration. Emerging work further indicates that microglial purinergic signalling operates within a coordinated receptor network in which ATP/ADP-mediated activation of P2Y receptors drives intracellular calcium dynamics that shape cytokine production, process motility, and microglia–neuron interactions [72]. In this context, P2Y12R-dependent Gi signalling may integrate with calcium-dependent pathways and other purinergic receptors to fine-tune inflammatory mediator release and injury sensing. However, the downstream signalling pathways linking P2Y12R activity to prolonged microglial activation remain to be defined [18, 73], and are likely to involve context-dependent interactions with additional purinergic, calcium-dependent, and neuroimmune signalling cascades.

Functionally, microglial P2Y12R signalling was associated with motor impairment across both acute and subchronic MPTP paradigms. Conditional receptor deletion mitigated MPTP-induced deficits in motor coordination and reduced abnormal increases in forelimb grip force, consistent with attenuation of rigidity-like behaviour rather than loss of muscle strength, a distinction that parallels clinical observations in Parkinson’s disease [74, 75]. Notably, the pattern of behavioural rescue differed between acute and subchronic models, suggesting that slow ramp-up deficits in the acute MPTP model may predominantly reflect nonspecific neurotoxic effects rather than progressive parkinsonian motor dysfunction, whereas microglial P2Y12R signalling contributes more prominently to bradykinetic and rigidity-associated motor impairments that persist during progressive dysfunction [76].

The subchronic MPTP experiments further demonstrate that, following resolution of the acute neurotoxic phase, activated microglia remain associated with a sustained inflammatory environment [18]. Although acute MPTP intoxication elicited a broad inflammatory response, the subchronic phase was characterized by a more restricted cytokine signature, with elevated TNFα and IL-1β selectively detected in the striatum. By contrast, IL-6 and IFN-γ levels remained unchanged, and the chemokines MCP-1 and KC were below the limit of detection in both the substantia nigra pars compacta and striatum. This pattern suggests a shift from the broad inflammatory response observed during acute intoxication toward a more limited, persistent inflammatory profile during the subchronic phase. Consistent with this interpretation, elevated TNFα, IL-1β, and IL-6 have been detected in the cerebrospinal fluid and peripheral blood of patients with Parkinson’s disease, indicating that elements of this cytokine profile are also observed in the human condition [77]. Within this context, interrupting microglial P2Y12R signalling during ongoing neurodegeneration remained beneficial, preserving dopaminergic neurons, dampening chronic microglial activation, and improving motor performance weeks after toxin exposure. Together, these findings are consistent with a role for microglial P2Y12R in maintaining subchronic inflammatory signalling and suggest that microglial pathways may remain therapeutically accessible during established stages of disease, a consideration of particular relevance for translational strategies in Parkinson’s disease.

To explore translational feasibility, we identified and pharmacologically characterized a novel class of small-molecule P2Y12R antagonists [78, 79]. Classical dose–response and Schild analyses confirmed that PSB0739 and ticagrelor produce predictable, surmountable inhibition of P2Y12R signalling *in vitro*, validating the assay system. Clopidogrel was not evaluated in this context because it is a prodrug requiring metabolic activation and is therefore inactive in cell-based assays [80], underscoring the necessity for direct-acting P2Y12R antagonists when interrogating central receptor function. Structure-based screening subsequently identified compounds 2170 and 2171, which inhibited P2Y12R signalling *in vitro* and were therefore advanced as first-generation pharmacological tools. Increasing concentrations produced rightward shifts of the 2-MeSADP concentration–response curves, yielding apparent pA₂ estimates of 6.05 and 6.7 for 2170 and 2171, respectively, although these profiles did not permit definitive classification as classical competitive or surmountable antagonists. Intracisternal administration of compound 2171, which bypasses the BBB, subsequently reproduced the neuroprotective phenotype associated with receptor blockade, providing pharmacological proof-of-concept for central P2Y12R engagement *in vivo*. Notably, these compounds were drawn from a focused library of N-substituted phenothiazines and related heterocyclic derivatives with known multidrug resistance (MDR) modulatory or cytotoxic activities, previously shown to inhibit ABC transporters such as P-glycoprotein, MRP1, and BCRP [79], providing a mechanistic rationale for potential central exposure despite their apolar physicochemical features. Phenothiazine derivatives are known to interact with multiple molecular targets; therefore, further medicinal chemistry optimization will be required to enhance receptor selectivity [81]. Accordingly, the observed neuroprotection should be interpreted as proof of principle that this chemical scaffold can engage central P2Y12R signalling *in vivo*, rather than as evidence of BBB permeability or translational readiness. Importantly, the absence of cytotoxic effects at pharmacologically active concentrations supports target-directed activity and establishes an initial safety benchmark.

## 5. Conclusion

In conclusion, the present study identifies microglial P2Y12R signalling as a functionally relevant contributor to neuroinflammation, dopaminergic vulnerability, and motor dysfunction in experimental Parkinsonism. By integrating inducible microglia-specific genetic deletion with first-generation pharmacological tools, we establish a framework for interrogating central P2Y12R signalling *in vivo*. These findings support continued medicinal chemistry optimization of P2Y12R antagonists and further investigation of microglial purinergic signalling as a potential therapeutic strategy for Parkinson’s disease.

## Author contributions

**András Iring:** Data Curation, Investigation, Methodology, Visualization, Formal Analysis, Writing – original draft, Conceptualization, Project Administration, Writing – review & editing. **Dóra J. Kiss:** Conceptualization, Methodology, Writing – review & editing. **Mária Baranyi:** Methodology, Data Curation, Writing – review & editing. **Zsuzsanna T. Gál:** Investigation, Validation, Writing – review & editing. **Emese Herczku:** Investigation, Formal Analysis, Validation, Writing – review & editing. **Pál Tod:** Investigation, Formal Analysis, Data Curation, Writing – review & editing. **Attila Egyed:** Methodology, Formal Analysis, Validation, Writing – review & editing. **Kornél Demeter:** Resources, Writing – review & editing. **Ivar von Kügelgen:** Resources, Writing – review & editing. **Long-Jun Wu:** Resources, Writing – review & editing. **György M. Keserű:** Conceptualization, Methodology, Supervision, Resources, Funding Acquisition, Writing – review & editing. **Beáta Sperlágh:** Conceptualization, Funding Acquisition, Supervision, Project Administration, Resources, Writing – review & editing.

## Acknowledgements

We acknowledge the Unit of Behavioral Studies at the HUN-REN Institute of Experimental Medicine for access to behavioural equipment and thank Dóra Ömböli for excellent technical assistance with behavioural experiments and data acquisition. We also acknowledge the Light Microscopy Center at the HUN-REN Institute of Experimental Medicine for access to the Nikon Eclipse Ti-E inverted microscope and C2 laser confocal system and thank Dr. Pál Vági for expert technical support with microscopy. We are additionally grateful to Nikolett Lénárt (Neuroimmunology Research Group, HUN-REN Institute of Experimental Medicine) for her valuable assistance during experimental work.

## Declaration of generative AI and AI-assisted technologies in the manuscript preparation process

Generative AI–assisted tools were used only for language editing and stylistic refinement of an author-prepared manuscript; all scientific content, analyses, and conclusions were conceived and written by the authors.

## Data availability

All data supporting the findings of this study are included in the published article and Supplemental Information. Any additional information required to reanalyse the data reported in this paper is available from the corresponding author upon request.

## Abbreviations

2-MeSADP: 2-methylthioadenosine diphosphate
ABC: ATP-binding cassette
ACSF: Artificial cerebrospinal fluid
ANOVA: analysis of variance
ARRIVE: Animal Research: Reporting of In Vivo Experiments
BBB: blood–brain barrier
BCRP: breast cancer resistance protein
cAMP: cyclic adenosine monophosphate
CD68: cluster of differentiation 68
CHO-hP2Y12R: CHO cells expressing human
P2Y12R CHO-K1: Chinese hamster ovary cells
CNS: central nervous system
CNS MPO: CNS multiparameter optimization
CRC: concentration–response curves
CreERT2: tamoxifen-inducible Cre recombinase
CX3CR1: C-X3-C motif chemokine receptor 1
DA: dopamine
DAB: 3,3′-diaminobenzidine
DOPAC: 3,4-dihydroxyphenylacetic acid
Gi inhibitory: G protein
HPLC: high-performance liquid chromatography
HVA: homovanillic acid
Iba1: ionized calcium-binding adapter molecule 1
IFN-γ: interferon gamma
iKO: inducible knockout
IL-1β: interleukin-1 beta
IL-6: interleukin-6
KC: keratinocyte-derived chemokine
LDH: lactate dehydrogenase
MCP-1: monocyte chemoattractant protein-1
MDR: multidrug resistance
MPP+: 1-methyl-4-phenylpyridinium
MPTP: 1-methyl-4-phenyl-1,2,3,6-tetrahydropyridine
MRP1: multidrug resistance–associated protein 1
NA: noradrenaline
P2ry12: P2Y12-receptor gene (mouse)
P2Y12R: P2Y12-receptor
pA2: negative logarithm of antagonist concentration producing a twofold shift
PAMPA: passive membrane permeability assessment
PBS: phosphate-buffered saline
PD: Parkinson’s disease
PFA: paraformaldehyde
SD: standard deviation
SNpc: substantia nigra pars compacta
TH: tyrosine hydroxylase
TH+: tyrosine hydroxylase–positive
TNFα: tumor necrosis factor alpha

