## Supplemental Table for "Targeting microglial P2Y12-receptor signalling attenuates neurodegeneration in experimental Parkinson’s disease"

### Supplementary tables

**Supplementary Table 1. Calculated BBB and CNS MPO scores of selected compounds**

| Compound ID | BBB Score | CNS MPO Score |
|---|---|---|
| 422 | 4.00 | 3.06 |
| 1921 | 3.95 | 2.61 |
| 2170 | 4.02 | 3.85 |
| 2171 | 4.07 | 3.95 |
| 2161 | 1.45 | 2.93 |

**Supplementary Table 2. Tamoxifen-induced recombination efficiency across experimental cohorts.**

| Induction date | Paradigm | Treated (*n*) | Excluded (*n*) | Included (*n*) | Recombination Efficiency (%) |
|---|---|---|---|---|---|
| 11 Jun 2024 | Acute | 3 | 0 | 3 | 100 |
| 08 Aug 2024 | Acute | 11 | 2 | 9 | 81.8 |
| 09 Sep 2024 | Acute | 10 | 2 | 8 | 80 |
| 07 Oct 2024 | Acute | 3 | 3 | 0 | 0 (cohort excluded) |
| 04 Nov 2024 | Acute | 6 | 1 | 5 | 83.3 |
| 13 Dec 2024 | Acute | 4 | 0 | 4 | 100 |
| 02 Jan 2025 | Acute | 11 | 0 | 11 | 100 |
| 13 Feb 2025 | Acute | 10 | 0 | 10 | 100 |
| 14 Jul 2025 | Acute | 10 | 2 | 8 | 80 |
| 18 Aug 2025 | Acute | 7 | 4 | 3 | 42.9 |
| 09 Sep 2025 | Acute | 12 | 8 | 4 | 33.3 |
| 09 Sep 2025 | Subchronic | 5 | 5 | 0 | 0 (cohort excluded) |
| 09 Sep 2025 | Subchronic | 7 | 2 | 5 | 71.4 |
| 11 Sep 2025 | Subchronic | 4 | 0 | 4 | 100 |
| 21 Oct 2025 | Subchronic | 4 | 3 | 1 | 25 |
| 25 Dec 2025 | Subchronic | 10 | 2 | 8 | 80 |
| 21 Jan 2026 | Subchronic | 10 | 0 | 10 | 100 |
| **Total** | **—** | **119** | **26** | **93** | **78.1** |

Animals were excluded when lack of recombination was detected in assays where verification was feasible (immunofluorescence or immunoblot). Molecular and neurochemical analyses were performed on cohorts in which recombination efficiency had been independently confirmed at the cohort level. Two cohorts lacking detectable recombination were excluded prior to analysis and replaced.
