## Supplemental Methods for "Targeting microglial P2Y12-receptor signalling attenuates neurodegeneration in experimental Parkinson’s disease"

### Supplementary Methods.

#### HPLC Purity Analysis

Compound purity was assessed by high-performance liquid chromatography (HPLC) using a Shimadzu LCMS-2020 system equipped with a Reprospher 100 C18 column (5 µm, 100 × 3 mm). Samples were eluted using a gradient system consisting of eluent A (0.1 % formic acid in water) and eluent B (0.1 % formic acid in acetonitrile) at a flow rate of 1.5 ml / min. The gradient profile was as follows: 0 % B at baseline; linear increase to 100 % B over 7 minutes; 100 % B maintained from 7–8 minutes; return to initial conditions from 8–9 minutes; re-equilibration until 10 minutes. Column temperature was maintained at 30 °C, and the injection volume was 1 µl. Purity was determined by integration of UV absorbance peaks monitored at 254 nm.

#### Supplementary Figures

**Supplementary Figure 1. Representative HPLC chromatograms of selected P2Y12R antagonists.** Chromatograms corresponding to compounds 422, 1921, 2161, 2170, and 2171 (A-E) are shown. Peak integration at 254 nm was used for purity determination.

**Supplementary Figure 1A. Compound 422.**

#### ==== Shimadzu LabSolutions Browser Report ====

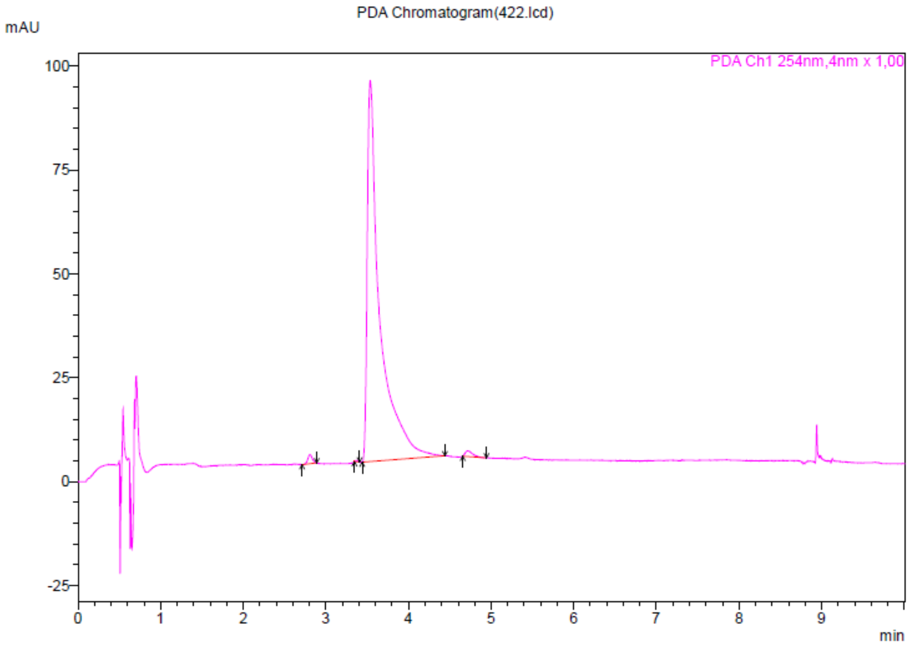


PDA Peak Table(422.lcd)

PDA Ch1 254nm

| Peak# | Ret. Time | Height | Area | Area% |
|---|---|---|---|---|
| 1 | 2,807 | 2155 | 8371 | 0,842 |
| 2 | 3,369 | 306 | 730 | 0,073 |
| 3 | 3,537 | 91300 | 976048 | 98,198 |
| 4 | 4,717 | 1416 | 8814 | 0,887 |
| Total | | 95178 | 993962 | 100,000 |

**Supplementary Figure 1A. Compound 1921.**

#### ==== Shimadzu LabSolutions Browser Report ====

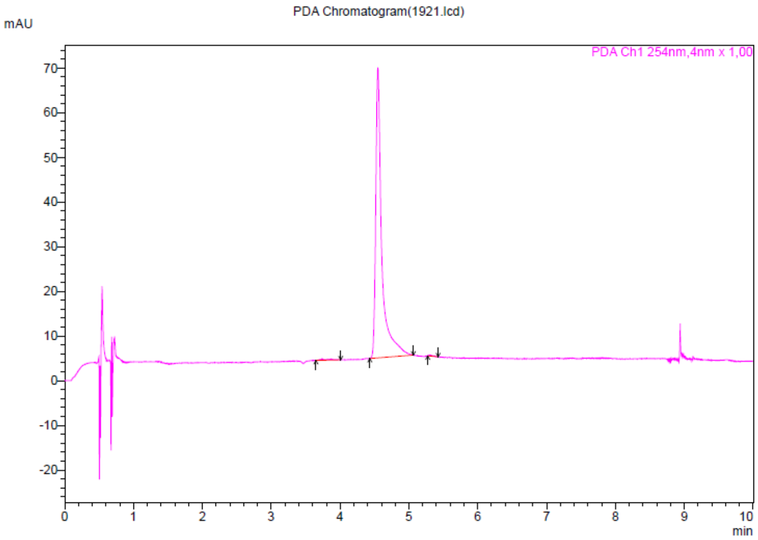


PDA Peak Table(1921.lcd)

PDA Ch1 254nm

| Peak# | Ret. Time | Height | Area | Area% |
|---|---|---|---|---|
| 1 | 3,745 | 257 | 2779 | 0,692 |
| 2 | 4,549 | 64744 | 397806 | 99,076 |
| 3 | 5,304 | 205 | 931 | 0,232 |
| Total | | 65206 | 401516 | 100,000 |

**Supplementary Figure 1C. Compound 2161.**

**==== Shimadzu LabSolutions Browser Report ====**

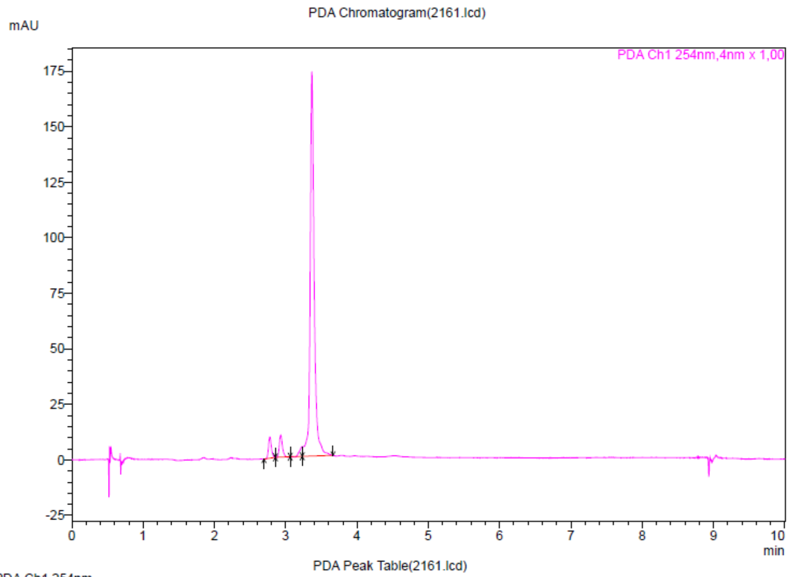


PDA Peak Table(2161.lcd)

PDA Ch1 254nm

| Peak# | Ret. Time | Height | Area | Area% |
|---|---|---|---|---|
| 1 | 2,777 | 9249 | 27059 | 3,584 |
| 2 | 2,929 | 9695 | 29510 | 3,908 |
| 3 | 3,225 | 4081 | 11057 | 1,464 |
| 4 | 3,367 | 171425 | 687413 | 91,043 |
| Total | | 194450 | 755040 | 100,000 |

**Supplementary Figure 1D. Compound 2170.**

**==== Shimadzu LabSolutions Browser Report ====**

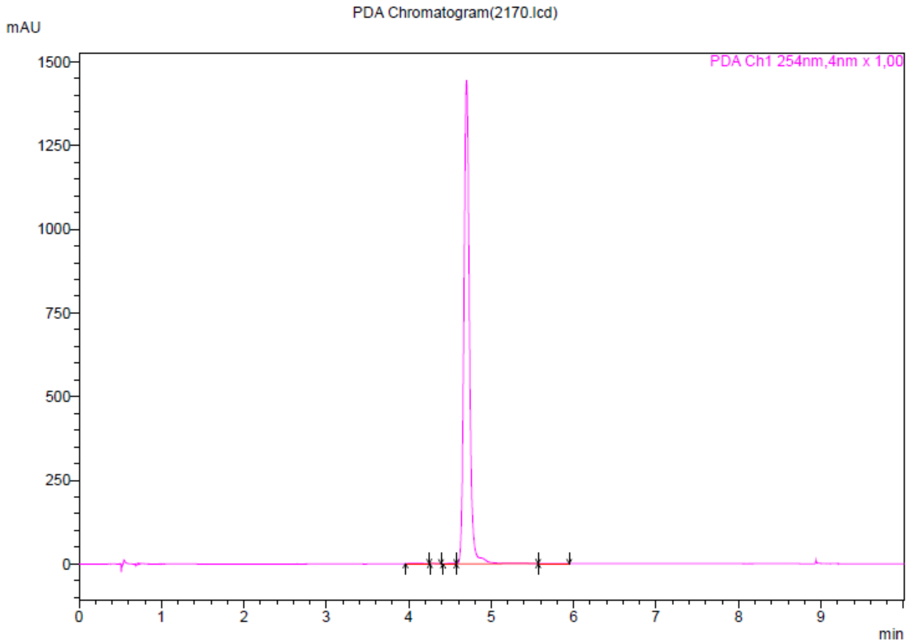


PDA Peak Table(2170.lcd)

PDA Ch1 254nm

| Peak# | Ret. Time | Height | Area | Area% |
|---|---|---|---|---|
| 1 | 4,057 | 1334 | 11876 | 0,176 |
| 2 | 4,330 | 317 | 975 | 0,014 |
| 3 | 4,506 | 1538 | 8196 | 0,121 |
| 4 | 4,701 | 1439738 | 6739943 | 99,636 |
| 5 | 5,581 | 347 | 3610 | 0,053 |
| Total | | 1443273 | 6764599 | 100,000 |

**Supplementary Figure 1E. Compound 2171.**

#### ==== Shimadzu LabSolutions Browser Report ====

PDA Chromatogram(2171.lcd)

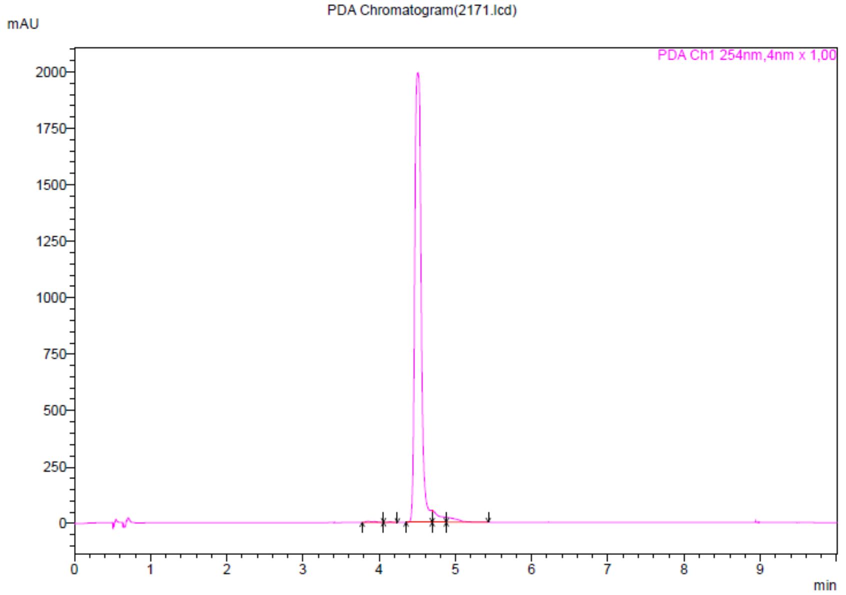


PDA Peak Table(2171.lcd)

PDA Ch1 254nm

| Peak# | Ret. Time | Height | Area | Area% |
|---|---|---|---|---|
| 1 | 3,853 | 4925 | 34220 | 0,278 |
| 2 | 4,146 | 2128 | 8281 | 0,067 |
| 3 | 4,506 | 1988568 | 11703974 | 95,217 |
| 4 | 4,697 | 52131 | 342353 | 2,785 |
| 5 | 4,881 | 22018 | 203084 | 1,652 |
| Total | | 2069769 | 12291911 | 100,000 |
